# Probing Allosteric Coupling of a Constitutively Open Mutant of the Ion Channel KcsA using Solid State NMR

**DOI:** 10.1101/567024

**Authors:** Zhiyu Sun, Yunyao Xu, Dongyu Zhang, Ann E McDermott

**Affiliations:** Department of Chemistry, Columbia University, New York, NY, 10027

## Abstract

Transmembrane allosteric coupling is a feature of many critical biological signaling events. Here we test whether transmembrane allosteric coupling controls the mean open time of the prototypical potassium channel KcsA in the context of C-type inactivation. Activation of KcsA is initiated by proton binding to the pH gate upon an intracellular drop in pH. Numerous studies have suggested that this proton binding also prompts a conformational switch leading to a loss of affinity for potassium ions at the selectivity filter and therefore to channel inactivation. We tested this mechanism for inactivation using a KcsA mutant (H25R/E118A) that has the pH gate open across a broad range of pH values. We present solid-state NMR measurements of this open mutant at neutral pH to probe the affinity for potassium at the selectivity filter. The potassium binding affinity in the selectivity filter of this mutant, 81 mM, is about 4 orders of magnitude weaker than that of wild type KcsA at neutral pH and is comparable to the value for wild type KcsA at low pH (pH ∼ 3.5). This result strongly supports our assertion that the open pH gate allosterically effects the potassium binding affinity of the selectivity filter. In this mutant the protonation state of a glutamate residue (E120) in the pH sensor is sensitive to potassium binding, suggesting that this mutant also has flexibility in the activation gate and is subject to transmembrane allostery.

**Significance statement:** Inactivation of potassium channels controls mean open times and provides exquisite control over biological processes. In the highly conserved C-type inactivation process, opening of the activation gate causes subsequent inactivation. We test whether the open state of the channel simply has a poor ability to bind the K^+^ ion. Previously, activated and inactivated states were stabilized using truncations or a significant pH drop. Here, we use the H25R/E118A constitutively open mutant of KcsA and also observe a large drop in potassium binding affinity. This provides strong evidence that channel opening causes an allosteric loss of ion affinity, and that the central feature of this universal channel inactivation process is loss of ion affinity at the selectivity filter.

## Introduction

The extracellular selectivity filter of potassium channels selectively allows potassium ions to cross biological membranes when activated at the intracellular gate by an external stimulus such as voltage or nucleotide binding. Generally, however, the channel spontaneously stops conducting even if potassium ions and activation stimuli are still present [1, 2]. This process typically occurs on a millisecond timescale after activation and is termed ‘C-type inactivation’. Inactivation is notably modulated by permeant ions and pore blockers [3]. Understanding the structural and mechanistic basis of C-type inactivation is expected to create insights into regulation of biomedically important channels such as the HERG channel that regulates the human heart and is significant for drug development [4, 5].

KcsA is a prototypical bacterial potassium channel which is pH-activated. The pH ‘gate’ is located at the intracellular face of the channel, formed by a group of key ionizable residues including H25, E118 and E120 [6–8]. Like its more complex eukaryotic homologs, KcsA undergoes inactivation on the millisecond timescale after activation by an intercellular pH drop [8, 9]. In previous studies, our group and others hypothesized that the site of inactivation is the selectivity filter, where potassium ions are bound and conducted through the membrane [1, 10–12]. This mechanism is termed activation-coupled inactivation and involves an allosteric coupling between the selectivity filter and activation gate (e.g. the pH gate in KcsA) that leads to a decrease in potassium affinity at the selectivity filter after activation. A direct measurement of the potassium ion affinity at the selectivity filter by solid state nuclear magnetic resonance (SSNMR) was carried out on KcsA embedded in a hydrated lipid environment [13]. The potassium ion affinity reduced upon acidification from to 4 ± 1 μM at pH 7.5 to K_app_ = 14 ± 1 mM at pH 3.5. Consequently, following activation of the channel at low pH, a reduction in potassium affinity at the selectivity filter is expected to lead to ion loss and conformational change at the selectivity filter, which in turn defeats efficient transmission. This is consistent with the electrophysiology studies showing that increasing extracellular potassium ion concentration could decrease the rate of inactivation [10, 11].

To test this model for allosteric coupling between the pH gate and the selectivity filter, we characterize a KcsA mutant that is constitutively open at neutral pH. pH gate residues H25, E118 and E120 have been suggested to form complex inter- and intra-subunit interactions at the cytoplasmic ends of the TM1 and TM2 transmembrane helices [14–16]. A triple mutant H25R/E120/E118A was shown to have the activation gate open across a broad range of pH values from 4 to 9 [17]. Separately, a solution NMR study also showed that H25 is key to the function of the pH gate [18]. Subsequently, it was shown only two mutations (to make the double mutant H25R /E118A) are sufficient to render the channel open at neutral pH [19]. Here we use the constitutively open H25R/E118A mutant to test whether the open channel indeed loses its ability to keep potassium ion loaded in the selectivity filter, and to control for nonspecific effects of the low pH used in other studies.

## Results

### The channel adopts similar conformations for the conductive and collapsed forms of the selectivity filter in the pH gate mutant as compared with wild type

NMR chemical shifts are precise indicators for protein structural and conformational changes. We conducted solid-state NMR experiments to investigate the structure of the H25R/E118A mutant of KcsA in a hydrated lipid bilayer environment (DOPE:DOPS = 9:1) with a 1:1 (w/w) protein to lipid ratio, as in our prior studies of wild type KcsA [13, 20–22]. The protein is well folded and displays high-quality NMR spectra at both the low (3.5) and high (7.5) pH values, and most chemical shift markers are similar to the corresponding markers for wild type (Figures S1 and S2). This confirms that the H25R/E118A mutant is well folded in lipid bilayers. Moreover, like the wild type, the H25R/E118A mutant displays two conformations at the selectivity filter. NMR chemical shifts report on the conformational change that occurs during K^+^ binding for the wild type, distinguishing the conductive and collapsed conformations seen in various crystal structures, as has been extensively discussed in prior literature. Both states can be observed at low pH and neutral pH, depending on potassium ion concentration. This indicates that the H25R/E118A mutant is well behaved and has the same conformational behavior at the selectivity filter as the wild type channel.

### Opening the pH gate significantly lowers potassium affinity at the selectivity filter at neutral pH

To test for the consequences of an open pH activation gate on function and binding in the selectivity filter in H25R/E118A, we measured the potassium affinity of this open pH gate mutant at neutral pH. We used nuclei in residues previously shown to have characteristic chemical shifts in the potassium binding site, including T75 (CA, CB, CG), T74 (CA, CB, CG) and V76 (CA, CG), to assess the populations of the apo and K^+^ bound states. The 2D 13C-^13^C DARR spectra of scaled chemical shift marker residues near the selectivity filter are shown in Figure 1 for various ambient potassium ion concentrations (at 10, 50, 75, 80, 85, 100, 150 mM [K^+^]). (Figure 1 a-d). The ionic strength was kept constant by compensating with sodium ions. As is the case for wild type KcsA, the K^+^ bound and apo state conformations do not exchange rapidly in this mutant, as evidences by the resolved peaks when the system is partially titrated [13].

**Fig. 1.**
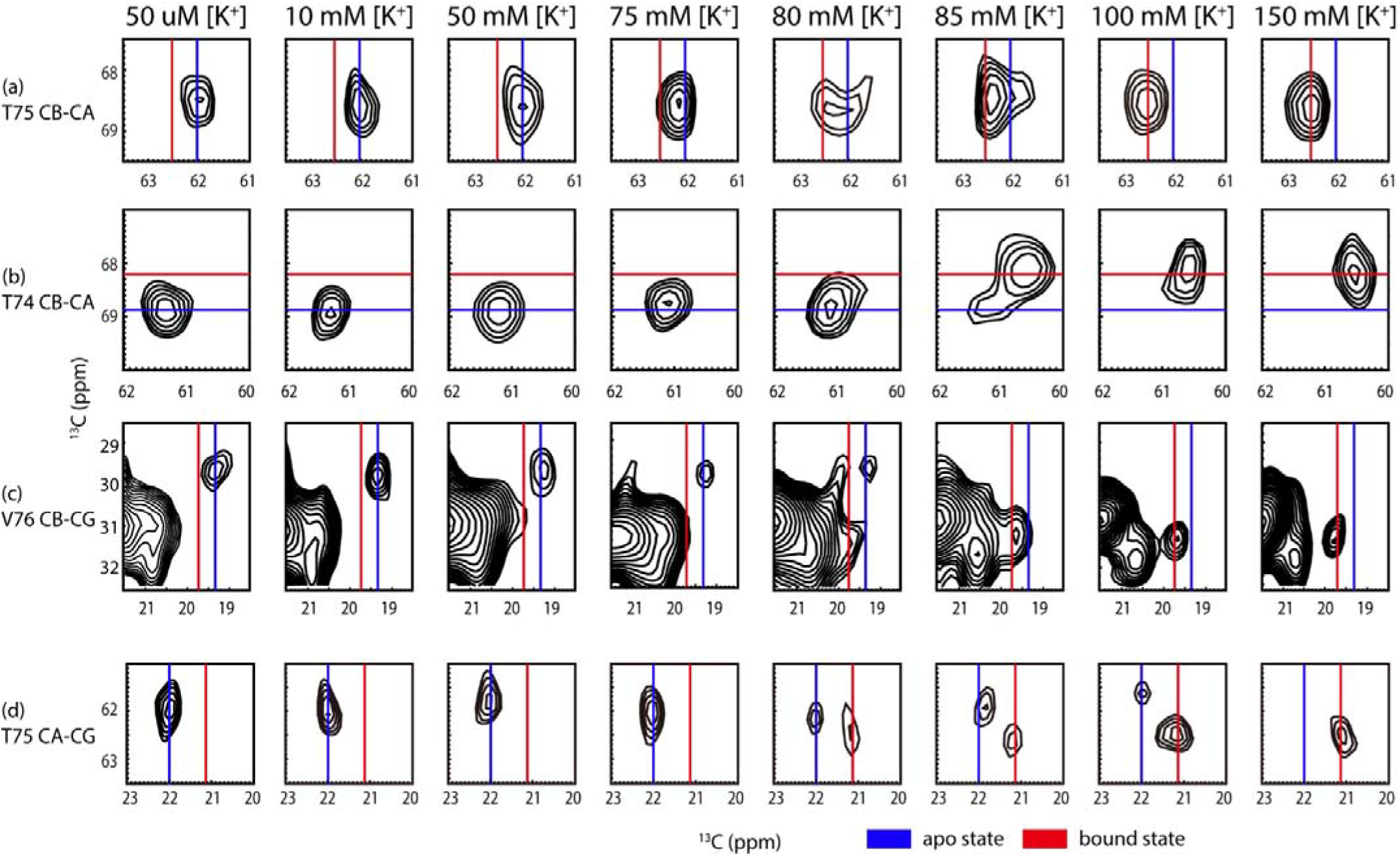
[K^+^] dependence of the intensities of the apo and bound state cross peaks of marker peaks near the selectivity filter in 2D ^13^C-^13^C (a-d) correlation spectra of H25R/E118A KcsA at pH 7.5. (a) T75 CB-CA. (b) T74 CB-CA. (c) V75 CB-CG. (d) T75 CA-CG. (a-d) show that the marker peaks shift from apo state in low [K^+^] to bound state in high [K^+^]. The contour level of the spectrum were set at 5 times noise level. Marker peaks could be integrated to calculate the population ratio of apo and bound states.

The ratio of potassium-bound KcsA to apo indicated by the NMR spectra is a function of ambient [K^+^]. This titration dependence was fit to a Hill model to extract K_app_ [13]. When the Hill coefficient was freely varied, K_app_ for K^+^ binding to H25R/E118A KcsA was best fit to 81 ± 1 mM, which is a similar value as for wild type KcsA at pH 3.5 (14 ± 1 mM) (Figure 2) [13]. These values are in stark contrast with the much tighter affinity for wild type KcsA at pH 7.5, 4 ± 1 μM. (Details of the Hill fit are shown in Table S1.) For the H25R/E118A mutant the transition from the bound to the apo state is steep as compared to that of the wild type KcsA at pH 3.5 and the best fit Hill coefficient was >8 in contrast to that for the wild type which was approximately 1. This can be explained as an increase in the cooperativity of the potassium binding process at the selectivity filter in this mutant as compared with the wild type. KcsA has multiple sites for proton binding and for potassium binding, so noncooperative binding is not necessarily expected. Differences in cooperatively with respect to the wild type may be related to the possibility that this mutant’s pH gate is open to a different degree than the wild type, or that it eliminates states with intermediate numbers of ions that occur for the wild type.

**Fig. 2.**
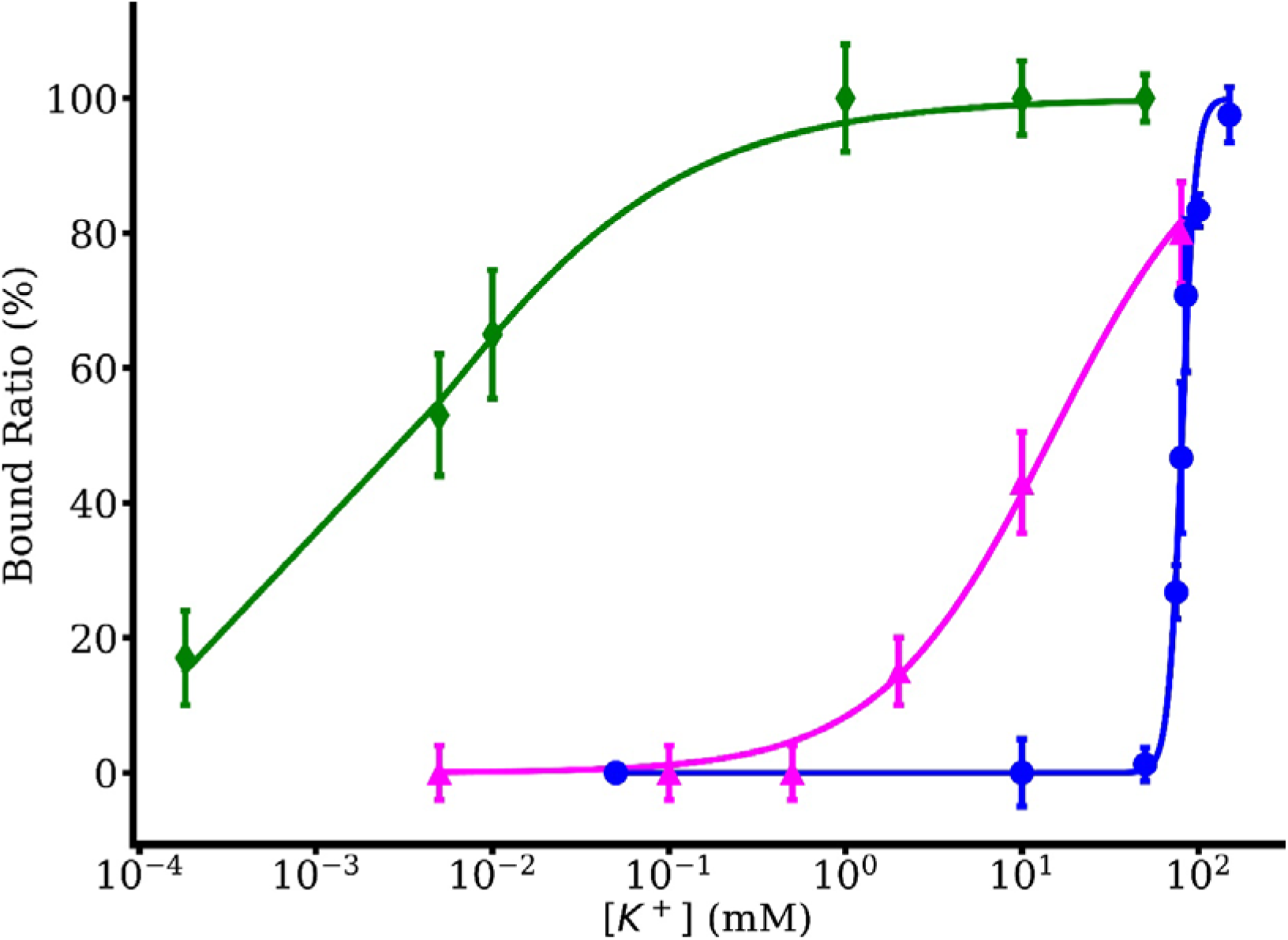
The bound ratio is plotted against [K^+^] to make the titration graph. The H25R/E118A (blue) is compared with our former data of wild type pH 7.5 (green) and wild type pH 3.5 (magenta). K_app_ of the open pH gate mutant was calculated to be 81 ± 1 mM by fitting the data to a Hill binding model compared with 4 ± 1 μM for wild type at pH 7.5 and 14 ± 1 mM for wild type at pH 3.5 [13]. Our data show that keeping the pH gate open with the H25R/E118A mutant further shifts the K^+^ affinity to a lower level.

As a control for nonspecific pH effects, the potassium affinity reduction, the affinity for this mutant at low pH was probed. At 80 mM ambient [K^+^], the fraction of H25R/E118A channels with K+ bound is approximately 40% both at low pH and neutral pH, suggesting that the K_app_ of the mutant is relatively pH insensitive (Figure S3) and that the nonspecific effects due to pH changes are not large.

In summary, these two very distinct perturbations (pH vs. mutation) appear to affect the thermodynamics at the selectivity filter very similarly, due to the fact that they both act to open the channel. This offers strong support for the hypothesis of allosteric activation coupled inactivation.

### The pH gate is coupled to ion binding in the constitutively open mutant

Allosteric coupling between the selectivity filter and the pH gate has been demonstrated in previous work [13, 23–28]. Not only does protonation and pH gate opening cause ion loss at the selectivity filter, but changes in ion occupation at the selectivity filter can reciprocally affect the protonation state of the pH gate residues E118 and E120 as well. For wild type KcsA, the peaks for E118 and E120 in ^13^C homonuclear SSNMR spectra are congested [13, 22]. Thus, we assign the peak at (34.11, 177.30) and (36.15, 183.10) (ppm) to CG-CD peak of residue E120 in the H25R/E118A mutant at high and low [K^+^] respectively, and we monitored the peak intensity changes as ambient [K^+^] was increased, as shown in Figure 3. Changes in the E120 CG-CD peak intensity and position indicates that this residue is protonated at low [K^+^], and deprotonated at high [K^+^], while the pH is held constant at a value of 7.5. At 50 μM [K^+^], E120 CG-CD appears to be essentially fully protonated, while at 80 mM [K^+^] the E120 CG-CD appears to be essentially fully deprotonated. Curiously, the titration curve for E120 is not identical to that of the selectivity filter markers, suggesting the existence of K^+^ ion sites of different affinity. These results show that in the H25R/E118A KcsA mutant, the pH gate and selectivity gate are allosterically coupled, as in to the coupling seen in the wild type [22].

**Fig. 3.**
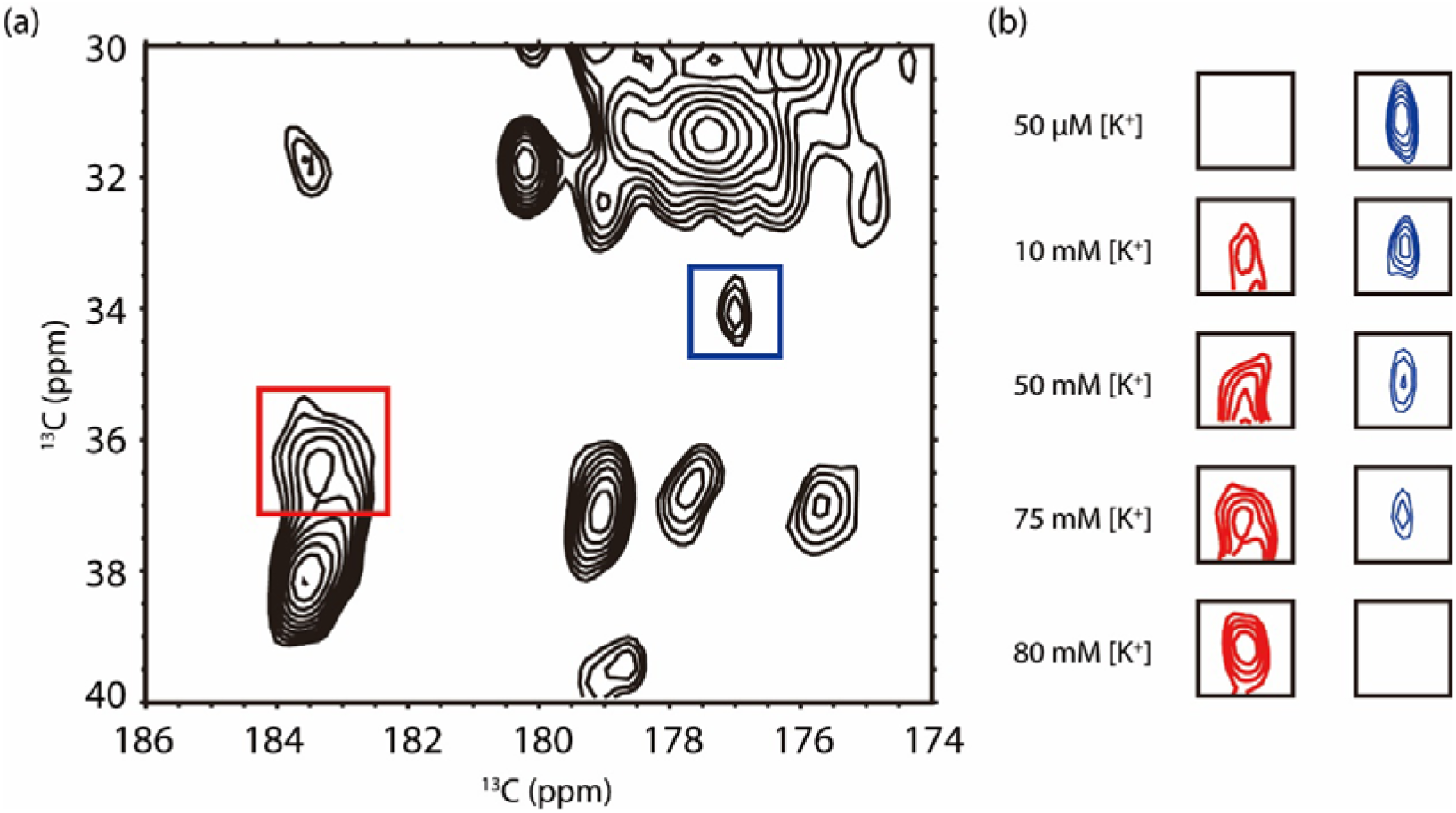
The effect of [K^+^] on E120 residue in the pH gate of the H25R/E118A KcsA mutant. 2D ^13^C-^13^C correlation spectrum of open pH gate KcsA at 75 mM [K^+^] is shown in (a). The peaks in the red and blue rectangles are the E120 CG-CD peaks in deprotonated and protonated state respectively. The protonated and deprotonated state change of the E120 peak across various [K^+^] is shown in (b). Protonated and deprotonated E120 CG-CD peaks at 50 μM, 10, 50, 75 and 80 mM are shown. At pH 7.5, E120 is protonated at low [K^+^], while at and above 80 mM [K^+^], E120 is completely deprotonated. In samples with [K^+^] larger than 80 mM, the E120 CG-CD peaks are all in the deprotonated state. This protonation state change provides clear evidence for pKa change of the glutamic acid residue.

### Electrophysiology of the H25R/E118A mutant

It was previously shown that although the H25R/E118A mutant is open, it does not conduct under standard electrophysiology conditions (pH 7.0, 100 mM [K^+^]), because it, like the wild type, inactivates when open. Inactivation can be suppressed by preparing the triple mutant E71A/ H25R/E118A [27]. The addition of the third mutation E71A has been shown to suppress inactivation in wild type and is commonly used as a “background” for many studies of KcsA to render it inactivationless, and make it more feasible to study the behavior of the activation gate in electrophysiology experiments [21, 29, 30]. Here, we also attempted another strategy to characterize the activation gate in electrophysiology experiments. We recently identified several essential participants in allosteric coupling between the activation and inactivation gate using NMR analysis [31], resulting in good agreement with residues identified in other molecular dynamics and electrophysiology studies [31, 32]. T74 is one of the key allosteric participants. The mutation T74S causes a strong reduction in allosteric coupling in KcsA at pH 5.0 and pH 3.5, and reduces the extent of inactivation dramatically.

Here we show that, the T74S/H25R/E118A mutant recovers activity (i.e. has reduced inactivation) allowing us to demonstrate that the activation gate is open. These results further support the hypothesis that the H25R/E118A mutant has an open pH gate at neutral pH. Also, they support our hypothesis that T74 is crucial for allosteric coupling between the pH gate and the selectivity filter. However, the activity is not completely recovered, in that the T74S/H25R/E118A triple mutant exhibits 30% open probability from pH 4 to 7, whereas in E71A the open probability is close to 100%. This distinction was not seen in the wild type at low pH; for that case, both E71A and T74S essentially abolished inactivation. Apparently when the channel is constitutively open (in the H25R/E118A) the T74S mutation reduces but does not fully attenuate allosteric coupling (Figure S4). The fact that T74S essentially abolishes allosteric coupling when paired with the wild type pH sensor, whereas the T74S/H25R/E118A triple mutant only partially attenuates the coupling suggests that these mutations in the pH gate cause subtle differences in the energetics of the selectivity filter compared to the fully protonated (open) wild type. For example, it is possible that lacking some of the restraints of the hydrogen bonding network present in the wild type, the H25R/E118A mutant may open to a greater degree and thereby encourage inactivation—although this interpretation is purely speculative in absence of further evidence and investigation.

## Discussion

We documented transmembrane allosteric coupling in the inactivation process of a potassium channel using an open pH gate KcsA mutant H25R/E118A in comparison with the wild type. Specifically, we tested the hypothesis that C-type inactivation involves collapse of the selectivity filter as an allosteric response of the opening of the pH gate [33, 34]. By comparing potassium dependent SSNMR data from the open pH gate KcsA mutant to that of the closed wild type, we tested whether the open channel loses its grip on the potassium ion.

The activated state of the channel is difficult stabilize because of the extreme pH required and because of the inactivation process [33]. Efforts to trap the activated state and the early intermediates of the C-type inactivation process have involved lowering the pH and/or significant mutations [21, 22, 25, 27, 30, 35, 36]. Although lowering the pH seems to be an intuitively correct method to trap the activated and the inactivated state, we had concerns about nonspecific effects of low pH (on both lipids and protein). At physiological conditions, lipids are critical to sustaining the structure and function of KcsA and are not entirely acid stable [37–40]. Moreover, one can imagine a variety of other nonspecific pH effects on the protein. Here, instead of changing the pH we conducted measurements on a mutant of KcsA with constitutively open pH gate at neutral pH conditions, and effectively eliminated this concern. We observed a reduction in potassium affinity at the selectivity filter for H25R/E118A KcsA by four orders of magnitude when is the channel is opened neutral pH. This confirms that opening the pH gate significantly affects the affinity of the potassium ion. Evidently the pH gate does not directly affect the bound and apo structures of the selectivity filter as indicated by the NMR spectra of the extracellular markers in and near the selectivity filter. Since the pH gate and the selectivity filter are almost 30 Å away from each other, such a dramatic effect is referred to as an allosteric coupling.

Our results are in contrast with experimental results from an isothermal titration calorimetry (ITC) study on the similar mutant H25R/E118A/E120A with truncated C-terminus in a detergent environment. Under these conditions the affinity for K^+^ at pH 8 is 0.13 mM, close to the apparent potassium affinity value 0.15 mM for wild type KcsA in their measurement [41]. We attribute this inconsistency with our work potentially to the usage of detergent in the ITC experiment, which could affect the stability and thermodynamics of the system, as previously indicated in studies of KcsA and other systems [1, 42]. Truncation of the C-terminus likely also alters the thermodynamic properties of the channel; in electrophysiology experiments, KcsA with a truncated C-terminus showed a more rapid and complete inactivation [33, 43]. By contrast, the solid state NMR measurements reported here involve full length KcsA in detergent free hydrated bilayers and are therefore more comparable to physiological conditions.

## Conclusions

The inactivation process of a potassium channel was explored by contrasting potassium binding in the open vs closed state, using SSNMR. The potassium affinity of the open pH mutant is of the same order of magnitude as the affinity of wild type KcsA channel under acidic conditions. Thus, we consider that both are good models for the open state and that the open state has a loose affinity for potassium ion. This result further confirms the hypothesis; in C-type inactivation, the open pH gate would lead to ion loss in the selectivity filter. The pKa of E120 in the intracellular pH gate was also shown to be influenced by potassium ion binding at the selectivity filter, providing additional evidence for an allosteric coupling network in KcsA.

## Materials and Methods

### Expression and Purification of ^13^C, ^15^N-labeled open pH gate KcsA

The open pH gate H25R/E118A KcsA was recombinantly expressed using a plasmid previously prepared by the Nimigean laboratory [27]. The protein expression and purification protocol were based on our former work with minor modifications [44]. 4 μl of the open pH gate KcsA plasmids was added into 50 μl JM83 competent cells in a culture test tube. The mixture was incubated on ice for 30 minutes, then heat-shocked for 70 seconds in a 42 °C water bath and incubated on ice for another 3 minutes. 1 mL of S.O.C recovery medium (Invitrogen) was added, and the cells were incubated at 37 °C for 1 hour. The transformed cells were plated on Luria broth (LB) agar plates containing 100 μg/ml ampicillin and incubated overnight at 37 °C. Single colonies of the transformed cells were picked and transferred to 4× 5 ml LB with 100 μg/ml ampicillin. The preculture was incubated at 37 °C, 250 rpm, when the OD_600_ reached 1.0, the precultures were transferred into 4× 1 L LB with 100 μg/ml ampicillin using the same incubation conditions. Once the OD_600_ reached 0.9, the cells were harvested via centrifugation at 4 °C, 8000 rpm for 15 minutes and resuspended in 1 L M9 medium with 3.0 g ^13^C-labeled D-glucose, 0.5 g ^15^NH_4_Cl and 100 μg/ml ampicillin. The cells were incubated at 37 °C, 250 rpm for 1 hour to recover. The protein expression was induced with anhydrotetracycline (Sigma), and incubated at 20 °C, 330 rpm overnight. The induced cells were harvested via centrifugation at 4 °C at 8000 rpm for 15 minutes, resuspended in 100 mM KCl, 50 mM Tris, 2 mM decyl maltoside (DM), pH 7.5 equilibrium buffer and lysed by French Press. The cell membranes were extracted with 30 mM DM at 4 °C overnight. The unlysed cells and membranes were pelleted via centrifugation at 4 °C at 15000 rpm for 1 hour. The protein was purified by nickel affinity column and eluted with 200 mM Imidazole. Imidazole concentration was reduced by buffer exchange with the equilibrium buffer, and the protein solution was concentrated for reconstitution.

10 mg 9:1 w/w DOPE and DOPS (Avanti) were mixed in a 10 ml test tube; the chloroform was evaporated by nitrogen gas flow, and the lipids were resuspended into 100 mM KCl, 50 mM Tris, 2 mM DM, 0.01 mM sodium azide, pH 7.5 buffer at concentration of 10 mg/ml. The ^13^C,^15^N-labeled open pH mutant KcsA was reconstituted into DOPE/DOPS liposome at 1:1 protein to liposome weight ratio. The mixture was dialyzed against a solution of 50 mM Tris, x mM KCl, (100-x) mM NaCl (to compensate ionic strength, except for the 150 mM KCl sample) at pH 7.5 overnight at room temperature, changing three times. The protein/liposome pellets were harvested, then frozen and thawed at – 80 °C and room temperature three times to remove bulk water. The final sample was reduced to a volume of approximately 30 μl by again carrying out three freeze-thaw cycles (−80 °C to room temperature) and and the sample was then packed into a 3.2 mm Bruker rotor, which contained approximately 10 mg mutant KcsA.

### Solid-State Nuclear Magnetic Resonance and data analysis

Magic-angle-spinning (MAS) solid-state NMR spectra were measured on Bruker 750 MHz (17.6 T) and 900 MHz (21.1 T) Avance spectrometers at the New York Structural Biology Center (NYSBC). The MAS rate was 14 kHz and the variable temperature set point was 264 K. Typical radiofrequency (rf) field strengths were 93-109 kHz for ^1^H and 50-62.5 kHz for ^13^C. ^13^C chemical shifts were referenced externally to the downfield adamantane CH_2_ resonance chemical shift at 40.48 ppm [45].

2D ^13^C-^13^C Dipolar Assisted Rotational Resonance (DARR) experiments were measured to obtain homonuclear ^13^C correlation spectra [46, 47]. The DARR mixing time was 15 ms. SPINAL64 decoupling on the ^1^H channel was 85-90 kHz during acquisition [48].

We calculated the population of the bound and apo states as previously [13], using the bound and apo state of marker peaks for T75 CB-CA, T74 CB-CA, T75 AB-CG and V76 CB-CB, quantifying the integrals using a *sum over box* method in *Sparky* [49]. Peaks were identified based on our former assignments [22, 49]. The bound population was computed 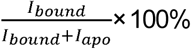. Normalized bound ratios of the protein for different ambient K^+^ concentrations were calculated by averaging the bound ratios of the marker peaks. Data were plotted and fit to a Hill equation in *OriginPro*: 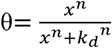, in which n is the Hill coefficient. The average bound population increased from 0% at 10 mM [K^+^] to 97% at 150 mM [K^+^]. In the range of 50 mM to 100 mM [K^+^], the major conformational transition takes place.

### Electrophysiology

T74S/H25R/E118A KcsA was prepared freshly using former protocols and purified with a gravity Nickel column and desalting column on FPLC. 5 mg DOPE:DOPG 3:1 (w/w) liposome were prepared as follows: lipids in chloroform were dried under nitrogen gas, and pentane was added to remove organic solvents; dry lipids were resuspended by sonication in 1 ml HEPES swelling buffer while slowly adding CHAPS detergent until clear. In total 30mg CHAPS was added. 2.5 ug T74S/H25R/E118A KcsA was added to the lipid/detergent solution, and the mixture was incubated at room temperature for 30 mins. A bio-bead column was used to remove the detergent, and the proteoliposomes were eluted at 8 ml and frozen in liquid nitrogen.

Electrophysiology experiments were carried out with a partition diameter of 100 μm. The lower chamber bath was at a pH of 4.0 (succinic acid buffer) and the upper chamber was at a pH of 7.0 (HEPES buffer, 100 mM K^+^). Data were collected in Clampex and processed in Clampfit.

## Acknowledgements

We thank Dr. Crina Nimigean of Cornell Weill University for assistance with the electrophysiology experiments. The NMR data were collected at the NYSBC with support from the Center on Macromolecular Dynamics by NMR Spectroscopy, a Biomedical Technology Research Resource supported by the NIH through Grant P41 GM118302. The NYSBC is also enabled by a grant from the Empire State Division of Science Technology and Innovation and by Office of Research Infrastructure Programs/NIH Facility Improvement Grant CO6RR015495. This work was supported by NIH Grant R01 GM088724 (to A.E.M).

## References

1. Cuello, L.G., et al., Structural basis for the coupling between activation and inactivation gates in K(+) channels. Nature, 2010. 466(7303): p. 272–5.

2. McCoy, J.G. and C.M. Nimigean, Structural correlates of selectivity and inactivation in potassium channels. Biochim Biophys Acta, 12012. 1818(2): p. 272–85.

3. Zachariae, U., et al., The molecular mechanism of toxin-induced conformational changes in a potassium channel: relation to C-type inactivation. Structure, 2008. 16(5): p. 747–54.

4. Alseikhan, B.A., et al., Engineered calmodulins reveal the unexpected eminence of Ca2+ channel inactivation in controlling heart excitation. Proc Natl Acad Sci U S A, 2002. 99(26): p. 17185–90.

5. Faber, E.S. and P. Sah, Ca2+-activated K+ (BK) channel inactivation contributes to spike broadening during repetitive firing in the rat lateral amygdala. J Physiol, 2003. 552(Pt 2): p. 483–97.

6. Cuello, L.G., et al., Proton-dependent gating in the Streptomyces K+ channel. Biophysical Journal, 1998. 74(2): p. A254–A254.

7. Heginbotham, L., et al., Single streptomyces lividans K(+) channels: functional asymmetries and sidedness of proton activation. J Gen Physiol, 1999. 114(4): p. 551–60.

8. Cortes, D.M., L.G. Cuello, and E. Perozo, Molecular architecture of full-length KcsA - Role of 1. cytoplasmic domains in ion permeation and activation gating. Journal of General Physiology, 2001. 117(2): p. 165–180.

9. Heginbotham, L., et al., Single Streptomyces lividans K+ channels: Functional asymmetries and sidedness of proton activation. Journal of General Physiology, 1999. 114(4): p. 551–559.

10. Bhate, M.P. and A.E. McDermott, Protonation state of E71 in KcsA and its role for channel collapse and inactivation. Proc Natl Acad Sci U S A, 2012. 109(38): p. 15265–70.

11. Wylie, B.J., M.P. Bhate, and A.E. McDermott, Transmembrane allosteric coupling of the gates in a potassium channel. Proc Natl Acad Sci U S A, 2014. 111(1): p. 185–90.

12. Linder, T., B.L. de Groot, and A. Stary-Weinzinger, Probing the energy landscape of activation gating of the bacterial potassium channel KcsA. PLoS Comput Biol, 2013. 9(5): p. e1003058.

13. Xu, Y., M.P. Bhate, and A.E. McDermott, Transmembrane allosteric energetics characterization for strong coupling between proton and potassium ion binding in the KcsA channel. Proc Natl Acad Sci U S A, 2017. 114(33): p. 8788–8793.

14. Miloshevsky, G.V. and P.C. Jordan, Open-state conformation of the KcsA K+ channel: Monte Carlo normal mode following simulations. Structure, 2007. 15(12): p. 1654–62.

15. Cuello, L.G., et al., A molecular mechanism for proton-dependent gating in KcsA. FEBS Lett, 2010. 584(6): p. 1126–32.

16. Cuello, L.G., et al., Design and characterization of a constitutively open KcsA. FEBS Lett, 2010. 584(6): p. 1133–8.

17. Thompson, A.N., et al., Molecular mechanism of pH sensing in KcsA potassium channels. Proc Natl Acad Sci U S A, 2008. 105(19): p. 6900–5.

18. Takeuchi, K., et al., Identification and characterization of the slowly exchanging pH-dependent conformational rearrangement in KcsA. J Biol Chem, 2007. 282(20): p. 15179–86.

19. Posson, D.J., et al., Molecular interactions involved in proton-dependent gating in KcsA potassium channels. J Gen Physiol, 2013. 142(6): p. 613–24.

20. Bhate, M.P., et al., Conformational Dynamics in the Selectivity Filter of KcsA in Response to Potassium Ion Concentration. Journal of Molecular Biology, 2010. 401(2): p. 155–166.

21. Bhate, M.P. and A.E. McDermott, Protonation state of E71 in KcsA and its role for channel collapse and inactivation. Proceedings of the National Academy of Sciences of the United States of America, 2012. 109(38): p. 15265–15270.

22. Wylie, B.J., M.P. Bhate, and A.E. McDermott, Transmembrane allosteric coupling of the gates in a potassium channel. Proceedings of the National Academy of Sciences of the United States of America, 2014. 111(1): p. 185–190.

23. Cuello, L.G., et al., A molecular mechanism for proton-dependent gating in KcsA. Febs Letters, 2010. 584(6): p. 1126–1132.

24. Cuello, L.G., et al., Structural basis for the coupling between activation and inactivation gates in K+ channels. Nature, 2010. 466(7303): p. 272–U154.

25. Imai, S., et al., Structural basis underlying the dual gate properties of KcsA. Proceedings of the National Academy of Sciences of the United States of America, 2010. 107(14): p. 6216–6221.

26. Thompson, A.N., et al., Molecular Interactions Involved in KCSA pH Gating. Biophysical Journal, 2011. 100(3): p. 273–273.

27. Posson, D.J., et al., Molecular interactions involved in proton-dependent gating in KcsA potassium channels. Journal of General Physiology, 2013. 142(6): p. 613–624.

28. Kim, D.M., et al., Conformational heterogeneity in closed and open states of the KcsA potassium channel in lipid bicelles. Journal of General Physiology, 2016. 148(2): p. 119–132.

29. Cordero-Morales, J.F., et al., Molecular determinants of gating at the potassium-channel selectivity filter. Nature Structural & Molecular Biology, 2006. 13(4): p. 311–318.

30. Thompson, A.N., et al., Molecular mechanism of pH sensing in KcsA potassium channels. Proceedings of the National Academy of Sciences of the United States of America, 2008. 105(19): p. 6900–6905.

31. Xu, Y., et al., Identifying coupled clusters of allostery participants through chemical shift perturbations. Proceedings of the National Academy of Sciences, 2019: p. 201811168.

32. Li, J., et al., Rapid constriction of the selectivity filter underlies C-type inactivation in the KcsA potassium channel. J Gen Physiol, 2018. 150(10): p. 1408–1420.

33. Cuello, L.G., et al., Structural mechanism of C-type inactivation in K+ channels. Nature, 2010. 466(7303): p. 203–U73.

34. Xu, Y.Y., M.P. Bhate, and A.E. McDermott, Transmembrane allosteric energetics characterization for strong coupling between proton and potassium ion binding in the KcsA channel. Proceedings of the National Academy of Sciences of the United States of America, 2017. 114(33): p. 8788–8793.

35. Cordero-Morales, J.F., L.G. Cuello, and E. Perozo, Voltage-dependent gating at the KcsA selectivity filter. Nature Structural & Molecular Biology, 2006. 13(4): p. 319–322.

36. Chakrapani, S., J.F. Cordero-Morales, and E. Perozo, A quantitative description of KcsA Gating I: Macroscopic currents. Journal of General Physiology, 2007. 130(5): p. 465–478.

37. Dart, C., Lipid microdomains and the regulation of ion channel function. Journal of Physiology-London, 2010. 588(17): p. 3169–3178.

38. Nakao, H., et al., pH-dependent promotion of phospholipid flip-flop by the KcsA potassium channel. Biochim Biophys Acta, 2015. 1848(1 Pt A): p. 145–50.

39. Schmidt, D., Q.X. Jiang, and R. MacKinnon, Phospholipids and the origin of cationic gating charges in voltage sensors. Nature, 2006. 444(7120): p. 775–9.

40. Valiyaveetil, F.I., Y.F. Zhou, and R. Mackinnon, Lipids in the structure, folding, and function of the KcsA K+ channel. Biochemistry, 2002. 41(35): p. 10771–10777.

41. Liu, S., et al., Ion-binding properties of a K+ channel selectivity filter in different conformations. Proc Natl Acad Sci U S A, 2015. 112(49): p. 15096–100.

42. Otzen, D.E., Protein unfolding in detergents: effect of micelle structure, ionic strength, pH, and temperature. Biophys J, 2002. 83(4): p. 2219–30.

43. Uysal, S., et al., Crystal structure of full-length KcsA in its closed conformation. Proceedings of the National Academy of Sciences of the United States of America, 2009. 106(16): p. 6644–6649.

44. Bhate, M.P., et al., Preparation of uniformly isotope labeled KcsA for solid state NMR: expression, purification, reconstitution into liposomes and functional assay. Protein Expr Purif, 2013. 91(2): p. 119–24.

45. Morcombe, C.R. and K.W. Zilm, Chemical shift referencing in MAS solid state NMR. J Magn Reson, 2003. 162(2): p. 479–86.

46. Takegoshi, K., S. Nakamura, and T. Terao, C-13-H-1 dipolar-driven C-13-C-13 recoupling without C-13 rf irradiation in nuclear magnetic resonance of rotating solids. Journal of Chemical Physics, 2003. 118(5): p. 2325–2341.

47. Takegoshi, K., S. Nakamura, and T. Terao, C-13-H-1 dipolar-assisted rotational resonance in magic-angle spinning NMR. Chemical Physics Letters, 2001. 344(5-6): p. 631–637.

48. Fung, B.M., A.K. Khitrin, and K. Ermolaev, An improved broadband decoupling sequence for liquid crystals and solids. Journal of Magnetic Resonance, 2000. 142(1): p. 97–101.

49. Goddard, T.D. and K. D.G., Sparky 3. University of California, San Francisco.

