## Supporting Information for "Probing Allosteric Coupling of a Constitutively Open Mutant of the Ion Channel KcsA using Solid State NMR"

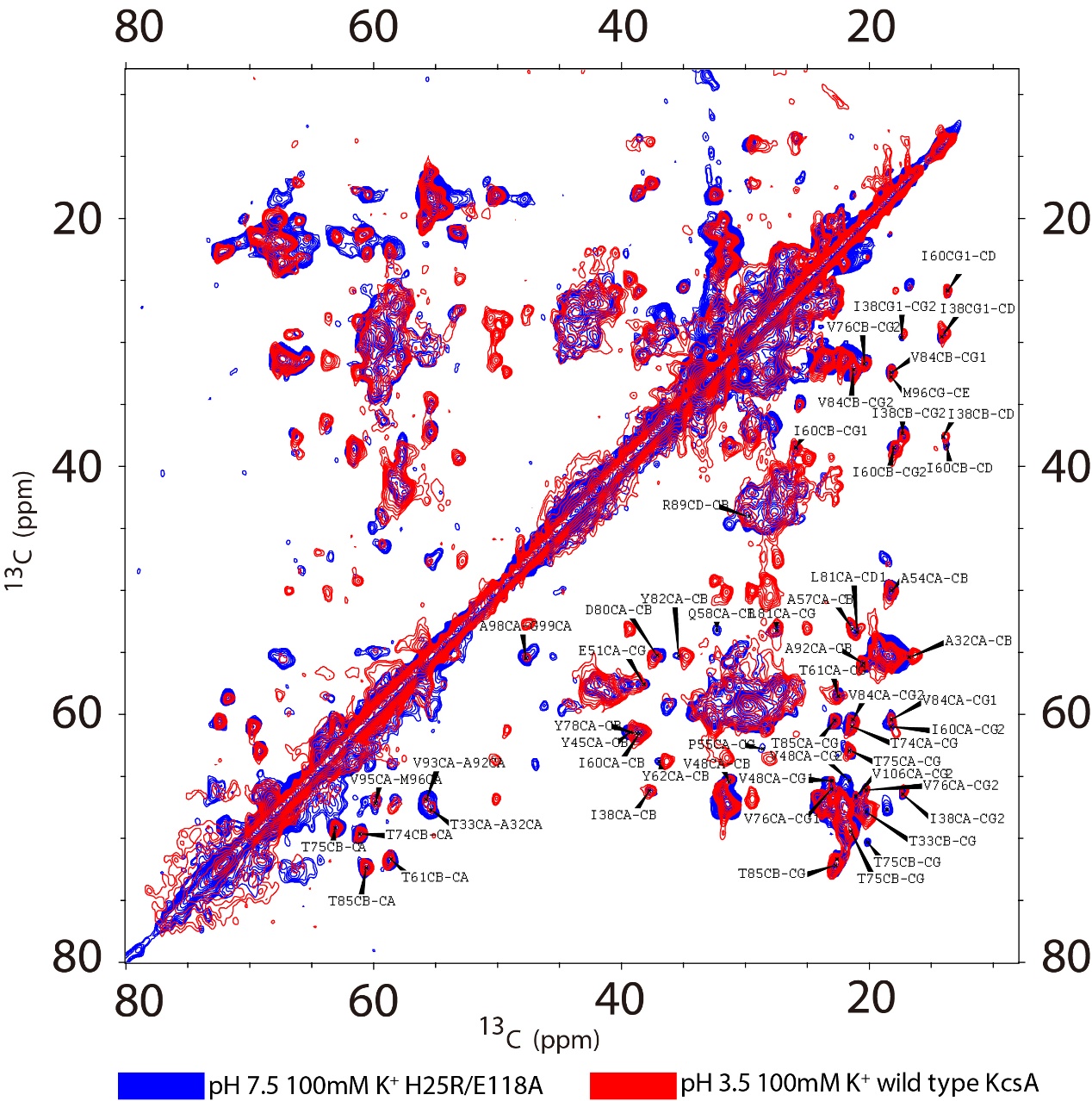

Fig. S1. 2D homonuclear ^13^C spectra of two samples that represent the open state of KcsA: H25R/E118A at pH 7.5 and 100mM K^+^ (blue) and wild type KcsA at pH 3.5 and 100 mM K^+^(red). The proline peaks are present in the wild type KcsA but not in the H25R/E118A mutant, because wild type KcsA is expressed Bl21(de3) cell line, however, the H25R/E118A mutant is expressed in JM83, a proline-auxotrophic cell line. The spectra these two models for the open state are largely identical.

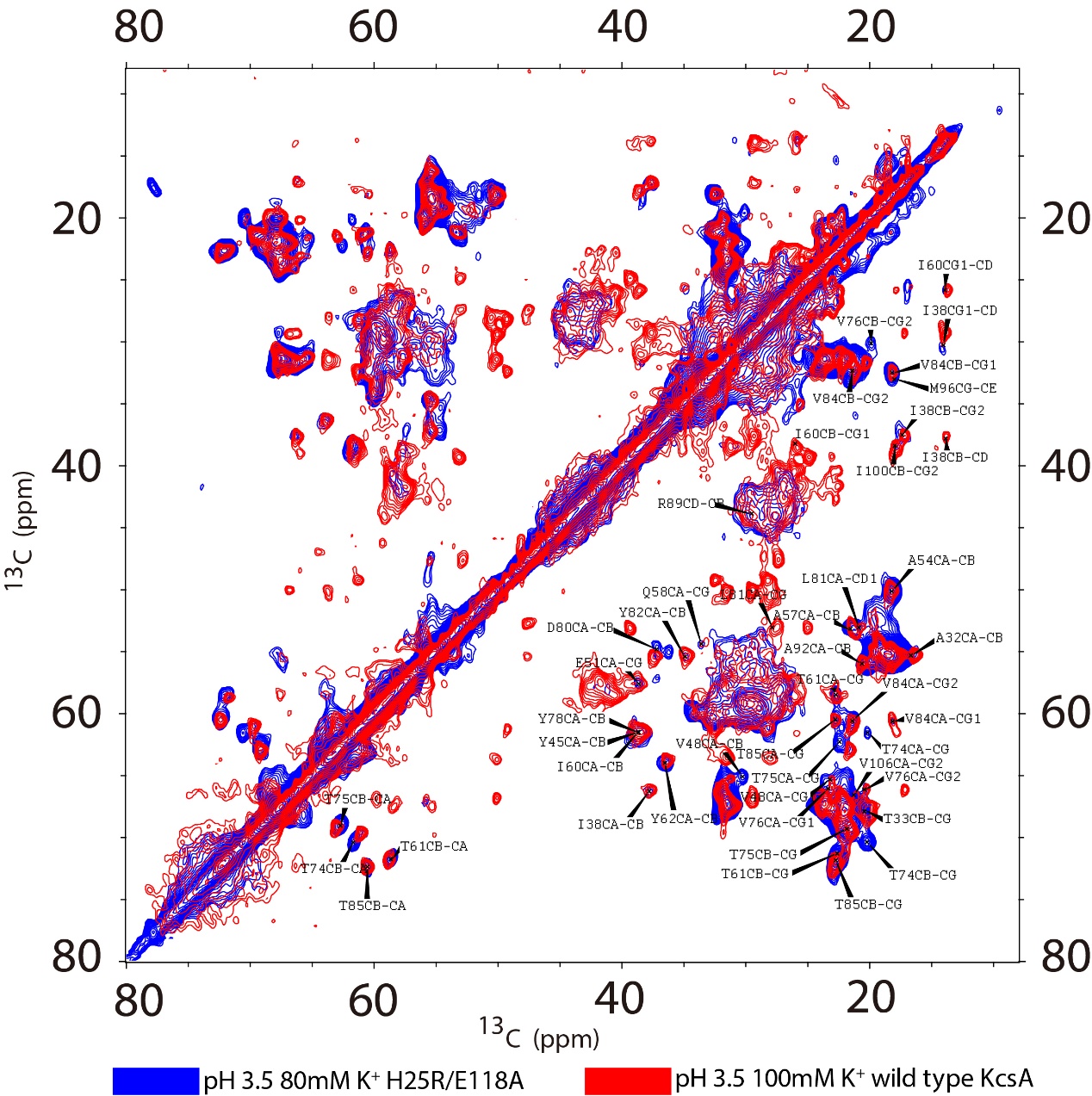

Fig. S2. 2D homonuclear ^13^C spectra of H25R/E118A at pH 3.5 and 80mM K^+^ and of wild type KcsA at pH 3.5 and 100 mM K^+^ are overlaid. The majority of the peaks of the mutant at low pH are identical to those of the wild type at the same pH (with the notable exception of the marker peaks in the selectivity filter since the mutant in this case is a mixed state). This indicates that the mutant is robust with respect to pH as is the wild type. Also the data suggests that the mutant has a somewhat weaker potassium binding affinity at pH 3.5 than at 7.5 (at this concentration the mutant be half titrated at pH 7.5, but favors the apo for pH 3.5).

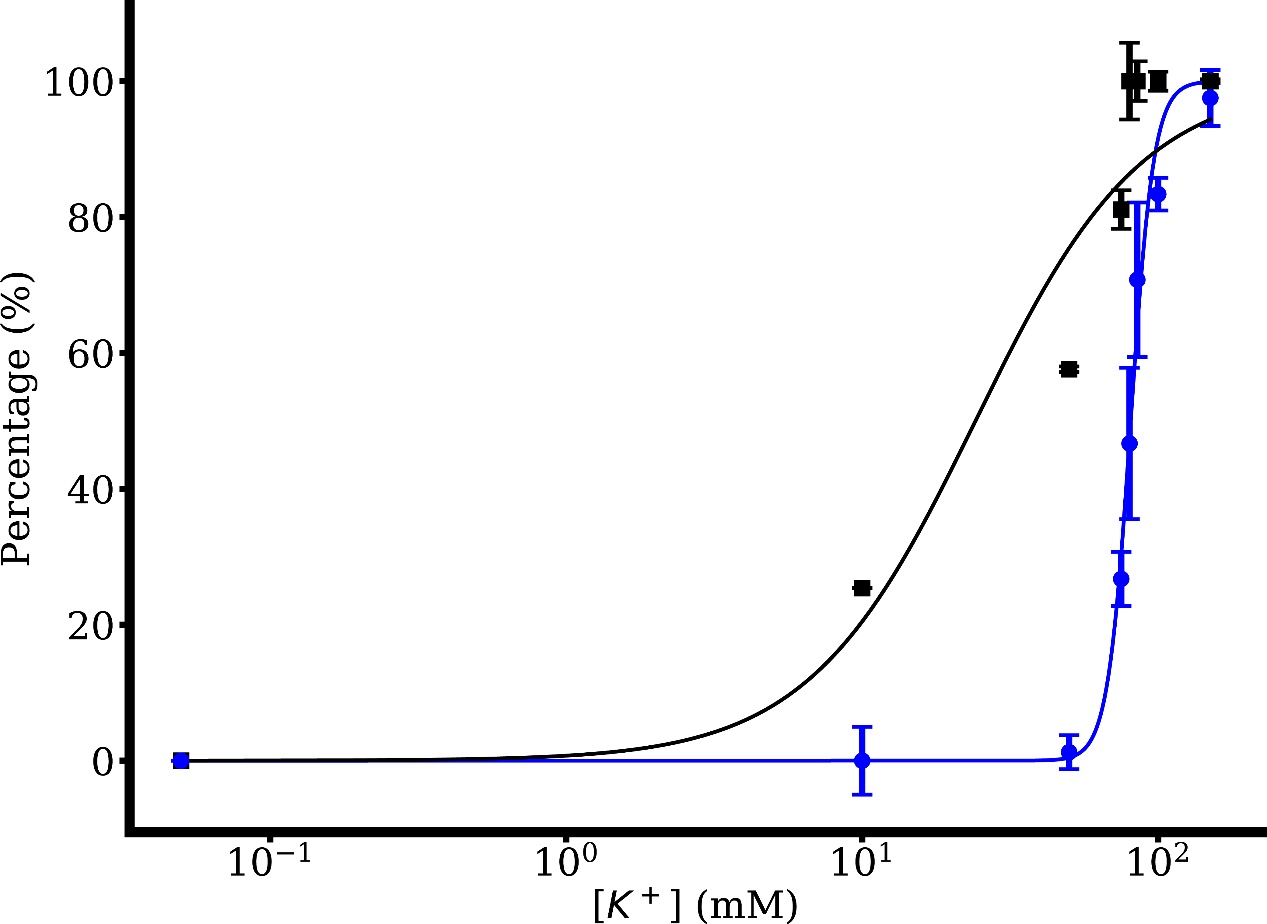

Fig. S3. The protonation state of E120 is monitored (percentage deprotonated state, black squares) in the same titration experiment where we probed binding at the selectivity filter (the percentage of the selectivity filter in the bound state, blue dots), both as a function of [K^+^]. The populations were assessed based on peak intensities as described in the materials and methods. The black curve is the fit of the E120 data to the Hill equation with K_app_ =24 ± 7 mM (R-Square = 0.91) and a Hill coefficient of n = 1.5. The titration behavior indicates that there is allosteric coupling between the intracellular and extracellular gates even in this constitutively open mutant. Compared to the selectivity filter, the pH sensor titrates at a slightly lower concentration, suggesting the presence of a distinct potassium binding sites.

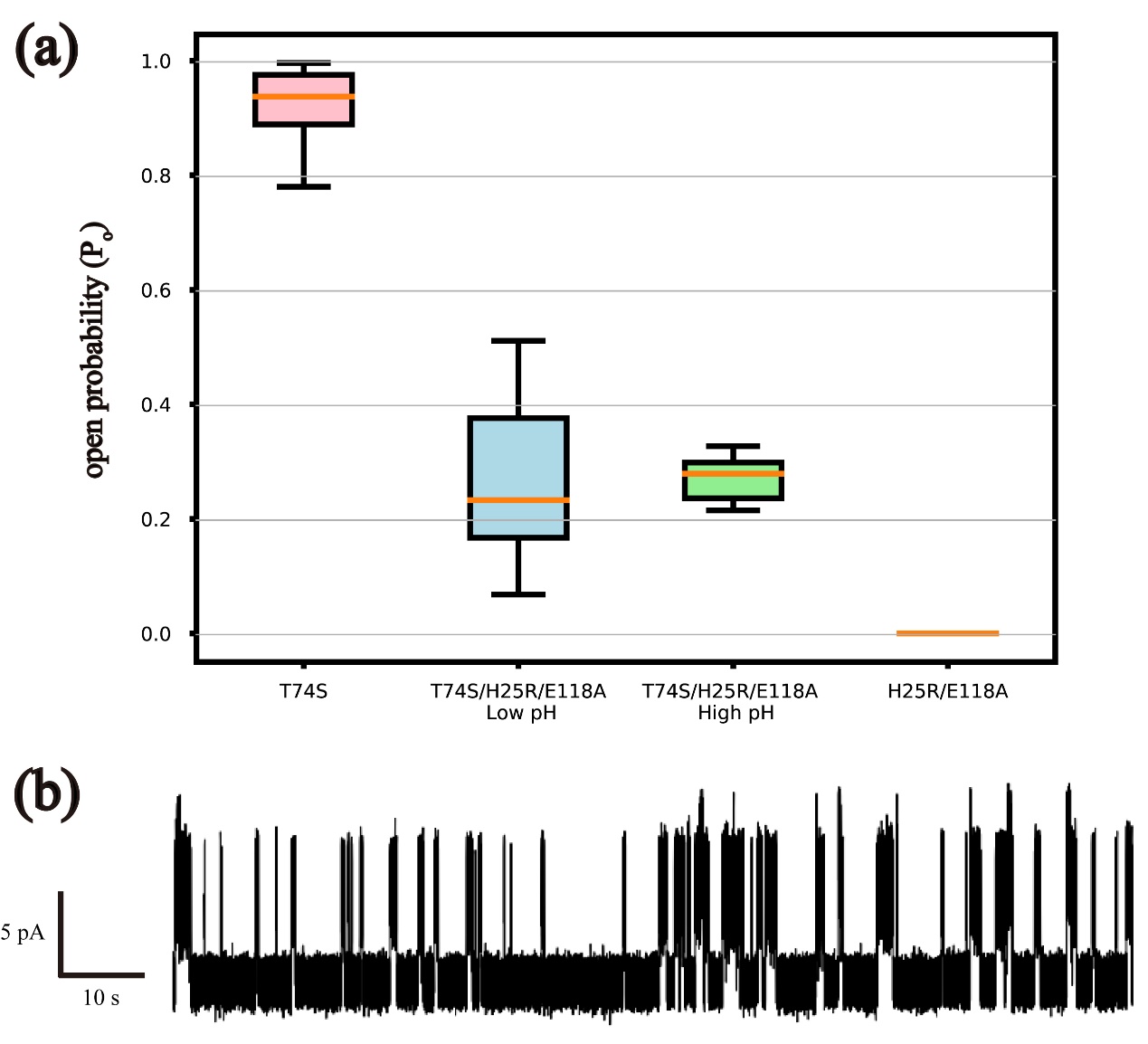

Fig. S4. Electrophysiology of open pH mutant and T74S triple mutant. (a). box plot of channel open probability. Open probability of T74S, H25RE118A_T74S low pH, H25RE118A_T74S high pH and H25RE118A are 90 ± 10%, 30 ± 20%, 27 ± 4% and 0 ± 0%. The upper chamber buffer pH of T74S and H25R/E118A is 7 and the lower chamber buffer pH of the two samples is 4. The pH of both T74S/H25R/E118A buffer in the upper chamber are 7, but the pH of bath buffer in the chamber are 4 and 7 corresponding to low and high pH. (b). A trace of single channel T74S/H25R/E118A at +100 mV. The channel is 30% open compared to the 0% Po of inactivated H25R/E118A, which suggests that the allosteric coupling between the dual gates are attenuated by the T74S mutation.

| model | Hill function | | | |
| --- | --- | --- | --- | --- |
| equation | y=$Vmax\frac{x^{n}}{x^{n}+{k_{d}}^{n}}$ | | | |
|  | (Vmax set to 1) | | | |
|  | k value (mM) | k std (mM) | Hill coefficient | adj r-square |
| WT pH7.5 | 0.0046 | 0.0009 | 1 | 0.967 |
| WT pH3.5 | 15 | 1 | 1 | 0.991 |
| H25RE118A pH7.5 | 100 | 60 | 1 | 0.533 |
| H25RE118A pH7.5 | 81 | 1 | 11 | 0.986 |

Table. S1. Parameters involved in optimized simulations to the Hill equation.

| Chemical shift markers of H25R/E118A at high [K+] and neutral pH | | | | | | | |
| --- | --- | --- | --- | --- | --- | --- | --- |
|  | CA | CB | CG | CG1 | CG2 | CD | CO |
| T74 | 61.05 | 69.65 | 21.41 | - | - | - | 176.3 |
| T75 | 63.03 | 69.63 | 21.08 | - | - | - | 172.4 |
| V76 | 66.06 | 31.73 | - | 23.04 | 20.31 | - | - |
| E120 | 59.40 | 36.31 | 34.11 | - | - | 177.30 | - |

Table. S2. Chemical shift markers of H25R/E118A at high [K+] and neutral pH

| Chemical shift markers of H25R/E118A at low [K+] and neutral pH | | | | | | | |
| --- | --- | --- | --- | --- | --- | --- | --- |
|  | CA | CB | CG | CG1 | CG2 | CD | CO |
| T74 | 61.18 | 69.94 | 20.92 | - | - | - | 177.2 |
| T75 | 62.89 | 69.54 | 21.27 | - | - | - | 172.4 |
| V76 | 66.06 | 31.33 | - | 23.11 | 20.27 | - | - |
| E120 | 59.40 | 36.31 | 36.15 | - | - | 183.10 | - |

Table. S3. Chemical shift markers of H25R/E118A at low [K+] and neutral pH

|  | CA | CB | CG/CG1 | CG2 | CD/CD1 | CE |
| --- | --- | --- | --- | --- | --- | --- |
| A32 | 55.37 | 16.76 | - | - | - | - |
| T33 | 67.72 | 67.87 | 20.24 | - | - | - |
| I38 | 66.31 | 37.58 | 29.57 | 17.34 | 14.10 | - |
| Y45 | 61.45 | 39.24 | - | - | - | - |
| V48 | 65.38 | 31.31 | 23.01 | 21.62 | - | - |
| E51 | 57.56 | - | 38.20 | - | - | - |
| A54 | 50.02 | 18.23 | - | - | - | - |
| P55 | 62.77 | - | 28.57 | - | - | - |
| A57 | 53.09 | 21.36 | - | - | - | - |
| Q58 | 53.21 | 32.29 | - | - | - | - |
| I60 | 61.26 | 38.46 | 25.88 | 18.13 | 13.70 | - |
| T61 | 58.57 | 71.68 | 22.52 | - | - | - |
| Y62 | 63.81 | 36.97 | - | - | - | - |
| T74 | 61.05 | 69.65 | 21.41 | - | - | - |
| T75 | 63.03 | 69.63 | 21.08 | - | - | - |
| V76 | 66.06 | 31.73 | 23.04 | 20.31 | - | - |
| Y78 | 61.51 | 38.78 | - | - | - | - |
| D80 | 55.25 | 37.13 | - | - | - | - |
| L81 | 53.28 | - | 27.50 | - | 20.98 | - |
| Y82 | 55.28 | 35.39 | - | - | - | - |
| V84 | 60.42 | 32.31 | 18.22 | 21.31 | - | - |
| T85 | 60.56 | 72.30 | 22.76 | - | - | - |
| A92 | 55.81 | 20.46 | - | - | - | - |
| M96 | 59.85 | - | 32.79 | - | - | 18.25 |

Table. S4. Assignment table of H25R/E118A at high [K+] and neutral pH

|  | CA | CB | CG/CG1 | CG2 | CD/CD1 | CE |
| --- | --- | --- | --- | --- | --- | --- |
| A32 | 55.36 | 16.74 | - | - | - | - |
| T33 | 67.72 | 67.87 | 20.29 | - | - | - |
| I38 | 66.31 | 37.61 | 29.67 | 17.34 | 14.07 | - |
| Y45 | 61.47 | 39.10 | - | - | - | - |
| V48 | 65.35 | 31.04 | 23.05 | 21.62 | - | - |
| E51 | 57.57 | - | 38.31 | - | - | - |
| A54 | 50.04 | 18.22 | - | - | - | - |
| P55 | 62.77 | - | 28.57 | - | - | - |
| A57 | 53.11 | 21.41 | - | - | - | - |
| Q58 | 53.51 | 32.29 | 33.59 | - | - | - |
| I60 | 61.29 | 38.46 | 25.89 | 18.13 | 13.73 | - |
| T61 | 58.57 | 71.61 | 22.58 | - | - | - |
| Y62 | 63.85 | 36.86 | - | - | - | - |
| T74 | 61.18 | 69.94 | 20.92 | - | - | - |
| T75 | 62.89 | 69.54 | 21.27 | - | - | - |
| V76 | 66.06 | 31.33 | 23.11 | 20.27 | - | - |
| Y78 | 61.51 | 38.75 | - | - | - | - |
| D80 | 55.13 | 37.15 | - | - | - | - |
| L81 | 53.22 | - | 27.58 | - | 20.95 | - |
| Y82 | 55.32 | 35.27 | - | - | - | - |
| V84 | 60.46 | 32.33 | 18.21 | 21.33 | - | - |
| T85 | 60.54 | 72.25 | 22.76 | - | - | - |
| A92 | 55.83 | 20.50 | - | - | - | - |
| M96 | 59.85 | - | 32.85 | - | - | 18.19 |

Table. S5. Assignment table of H25R/E118A at low [K+] and neutral pH

|  | CA | CB | CG/CG1 | CG2 | CD/CD1 | CE |
| --- | --- | --- | --- | --- | --- | --- |
| A32 | 55.36 | 16.74 | - | - | - | - |
| T33 | 67.72 | 67.87 | 20.29 | - | - | - |
| I38 | 66.31 | 37.61 | 29.67 | 17.34 | 14.07 | - |
| Y45 | 61.47 | 39.10 | - | - | - | - |
| V48 | 65.35 | 31.04 | 23.05 | 21.62 | - | - |
| E51 | 57.58 | - | 38.30 | - | - | - |
| A54 | 50.04 | 18.22 | - | - | - | - |
| P55 | 62.77 | - | 28.57 | - | - | - |
| A57 | 53.11 | 21.41 | - | - | - | - |
| Q58 | 53.51 | 32.29 | 33.59 | - | - | - |
| I60 | 61.29 | 38.46 | 25.89 | 18.13 | 13.73 | - |
| T61 | 58.57 | 71.61 | 22.58 | - | - | - |
| Y62 | 63.85 | 36.86 | - | - | - | - |
| T74 | 61.18 | 69.94 | 20.92 | - | - | - |
| T75 | 62.89 | 69.54 | 21.27 | - | - | - |
| V76 | 66.06 | 31.33 | 23.11 | 20.27 | - | - |
| Y78 | 61.51 | 38.75 | - | - | - | - |
| D80 | 55.13 | 37.15 | - | - | - | - |
| L81 | 53.22 | - | 27.58 | - | 20.95 | - |
| Y82 | 55.32 | 35.27 | - | - | - | - |
| V84 | 60.46 | 32.33 | 18.21 | 21.33 | - | - |
| T85 | 60.54 | 72.25 | 22.76 | - | - | - |
| A92 | 55.83 | 20.50 | - | - | - | - |
| M96 | 59.85 | - | 32.85 | - | - | 18.19 |

Table. S6. Assignment table of wild type KcsA at high [K+] and low pH
